# prewas: Data pre-processing for more informative bacterial GWAS

**DOI:** 10.1101/2019.12.20.873158

**Authors:** Katie Saund, Zena Lapp, Stephanie N. Thiede, Ali Pirani, Evan S. Snitkin

## Abstract

While variant identification pipelines are becoming increasingly standardized, less attention has been paid to the pre-processing of variants prior to their use in bacterial genome-wide association studies (bGWAS). Three nuances of variant pre-processing that impact downstream identification of genetic associations include the separation of variants at multiallelic sites, separation of variants in overlapping genes, and referencing of variants relative to ancestral alleles. Here we demonstrate the importance of these variant pre-processing steps on diverse bacterial genomic datasets and present prewas, an R package, that standardizes the pre-processing of multiallelic sites, overlapping genes, and reference alleles before bGWAS. This package facilitates improved reproducibility and interpretability of bGWAS results. Prewas enables users to extract maximal information from bGWAS by implementing multi-line representation for multiallelic sites and variants in overlapping genes. Prewas outputs a binary SNP matrix that can be used for SNP-based bGWAS and will prevent the masking of minor alleles during bGWAS analysis. The optional binary gene matrix output can be used for gene-based bGWAS which will enable users to maximize the power and evolutionary interpretability of their bGWAS studies. Prewas is available for download from GitHub.

**DATA SUMMARY:**

1. prewas is available from GitHub under the MIT License (URL: https://github.com/Snitkin-Lab-Umich/prewas) and can be installed using the command

~~~
devtools::install_github(“Snitkin-Lab-Umich/prewas”)
~~~
2. Code to perform analyses is available from GitHub under the MIT License (URL: https://github.com/Snitkin-Lab-Umich/prewas_manuscript_analysis)
3. All genomes are publicly available on NCBI (see Table S1 for more details)

**IMPACT STATEMENT:** In between variant calling and performing bacterial genome-wide association studies (bGWAS) there are many decisions regarding processing of variants that have the potential to impact bGWAS results. We discuss the benefits and drawbacks of various variant pre-processing decisions and present the R package prewas to standardize single nucleotide polymorphism (SNP) pre-processing, specifically to incorporate multiallelic sites and prepare the data for gene-based analyses. We demonstrate the importance of these considerations by highlighting the prevalence of multiallelic sites and SNPs in overlapping genes within diverse bacterial genomes and the impact of reference allele choice on gene-based analyses.

## INTRODUCTION

Bacterial genome-wide association studies (bGWAS) are frequently used to identify genetic variants associated with variation in microbial phenotypes such as antibiotic resistance, host specificity, and virulence (1–4). bGWAS methods can be classified into two general categories: those that use k-length nucleotide sequences (kmers) as features (e.g. (3,5–7)), and those that use defined variant classes such as single nucleotide polymorphisms (SNPs), gene presence/absence, or insertions/deletions (indels) as features (e.g. 4,8–12). bGWAS can be performed using individual variants or by grouping variants into genes or pathways (i.e. performing a burden test). While there have been efforts to standardize variant identification protocols (13,14), less attention has been paid to the downstream processing of variants prior to their use for applications like bGWAS. In this paper, we focus on pre-processing of SNPs (Figure 1A); however, the ideas and methods we discuss with respect to SNPs can be extended to other genetic variants.

**Figure 1:**
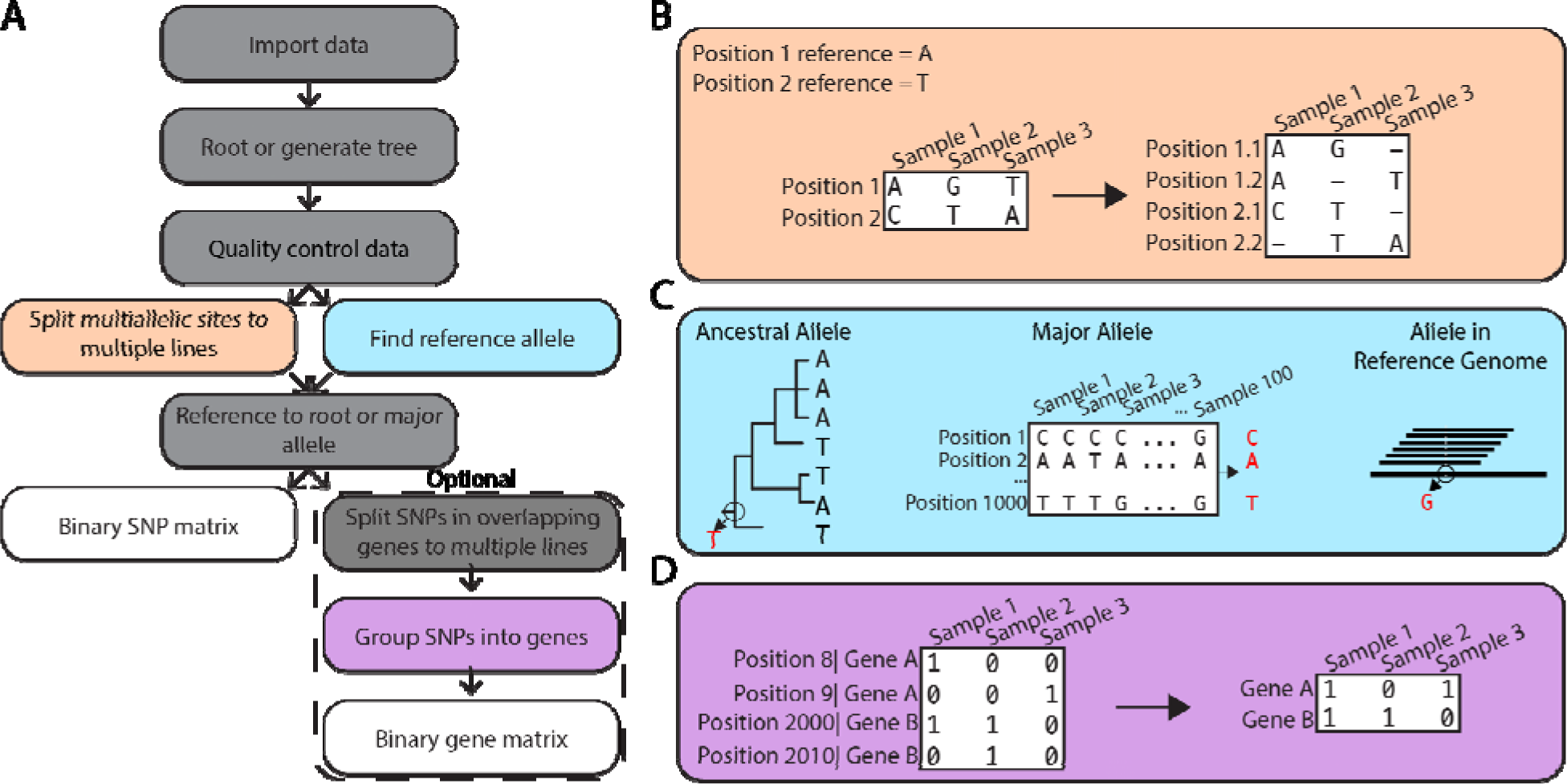
prewas workflow. (A) Overview of the prewas workflow. Grey and colored boxes: processing steps. White boxes: output generated. (B) Multi-line representation of multiallelic sites. (C) Possible methods to find a reference allele. The ancestral allele method and the major allele method are implemented in prewas. (D) Grouping SNPs into genes.

One aspect of pre-processing for SNP-based bGWAS is handling multiallelic sites. A site in the genome is considered multiallelic when more than two alleles are present at that locus (Figure 1B). Multiallelic sites do not fit neatly into the framework of most bGWAS methods, which often require a binary input (e.g. 3,4). Furthermore, the alternative minor alleles at a single site may impact the encoded protein to different extents, and therefore considering them separately may allow users to uncover otherwise masked relationships between genotype and phenotype.

Grouping SNPs by genes or metabolic pathways (Figure 1D) prior to performing bGWAS increases power and reduces collinearity (3,15,16). When performing gene-based analyses, two pre-processing steps may include choosing a reference allele for each SNP (Figure 1C) and assigning SNPs in overlapping gene pairs. The reference allele is the nucleotide relative to which variants are defined. Choice of reference allele is particularly important when grouping SNPs by gene to ensure that the direction of evolution for each SNP is preserved. Additionally, overlapping genes are common in bacteria (17,18). SNPs shared by overlapping gene pairs may be assigned to both genes in a gene-based analysis.

To determine the importance of variant pre-processing methods for bGWAS, we investigated the prevalence of multiallelic sites, mismatches in reference allele choice, and SNPs in overlapping genes in 9 bacterial datasets. Our analysis indicates that multiallelic sites are common in large, diverse bacterial datasets, there are frequently mismatches between different reference allele choices, and SNPs in overlapping genes often have discordant functional impacts. Therefore, pre-processing decisions have the potential to impact to bGWAS results.

We implemented a solution in the R package prewas to handle the nuances of variant preprocessing to enable more robust and reproducible bGWAS analyses (Figure S1). The output of prewas can be directly input into bGWAS tools that require a binary matrix as an input (e.g. (3,4)). Prewas can be downloaded from GitHub.

## METHODS

### Datasets

The collection of datasets we used for data analysis and the corresponding bioprojects are listed in Table S1 (19–30). All of these datasets contain whole-genome sequences of the bacterial isolates.

### Variant calling & tree building

SNP calling and phylogenetic tree reconstruction were performed on each dataset as described in (23). The variant calling pipeline can be found on GitHub (https://github.com/Snitkin-Lab-Umich/variant_calling_pipeline). In short, variant calling was performed with samtools v0.1.18 (31) using the reference genomes listed in Table S1, and trees were built using IQ-TREE v1.5.5 (32).

### Functional impact prediction

The functional impact of each SNP was predicted using SnpEff (33). Variants are categorized by SnpEff as low impact (e.g. synonymous mutations), moderate impact (e.g. nonsynonymous mutations), or high impact (e.g. nonsense mutations). Only variants in coding regions were included in analyses.

### Data analysis

Statistical analyses and modeling were conducted in R v3.6.1. The analysis code and data are available at: github.com/Snitkin-Lab-Umich/prewas_manuscript_analysis. The R packages we used can be found in the prewas.yaml file on GitHub (github.com/Snitkin-Lab-Umich/prewas; 34–43), and can be installed using miniconda (44).

#### Multiallelic sites

Linear regressions were modeled with percentage of variants that are multiallelic as the response variable and either number of samples or mean pairwise SNP distance as the predictor. R^2^ values are reported.

#### Reference alleles

For each dataset, the reference genome allele, major allele, and ancestral allele were identified and the number of mismatches between them was quantified. Ancestral reconstruction was performed in R using the ape::ace function with ape v5.3 (34).

#### Allele convergence

We recorded the number of times each allele arises on the tree, as inferred from ancestral reconstruction, and then subtracted 1 to calculate the number of convergence events for each allele.

## RESULTS & DISCUSSION

To maximize the potential for identifying genetic variation associated with a given phenotype using bGWAS, care must be taken in the pre-processing stage. Here we focus on three aspects of variant pre-processing and evaluate their potential downstream importance for bGWAS analysis. In particular, we report on the prevalence of multiallelic sites, mismatches between reference allele choice, and variants in overlapping genes across 9 bacterial datasets from various species and of varying genetic diversity (Table 1).

**Table 1:**
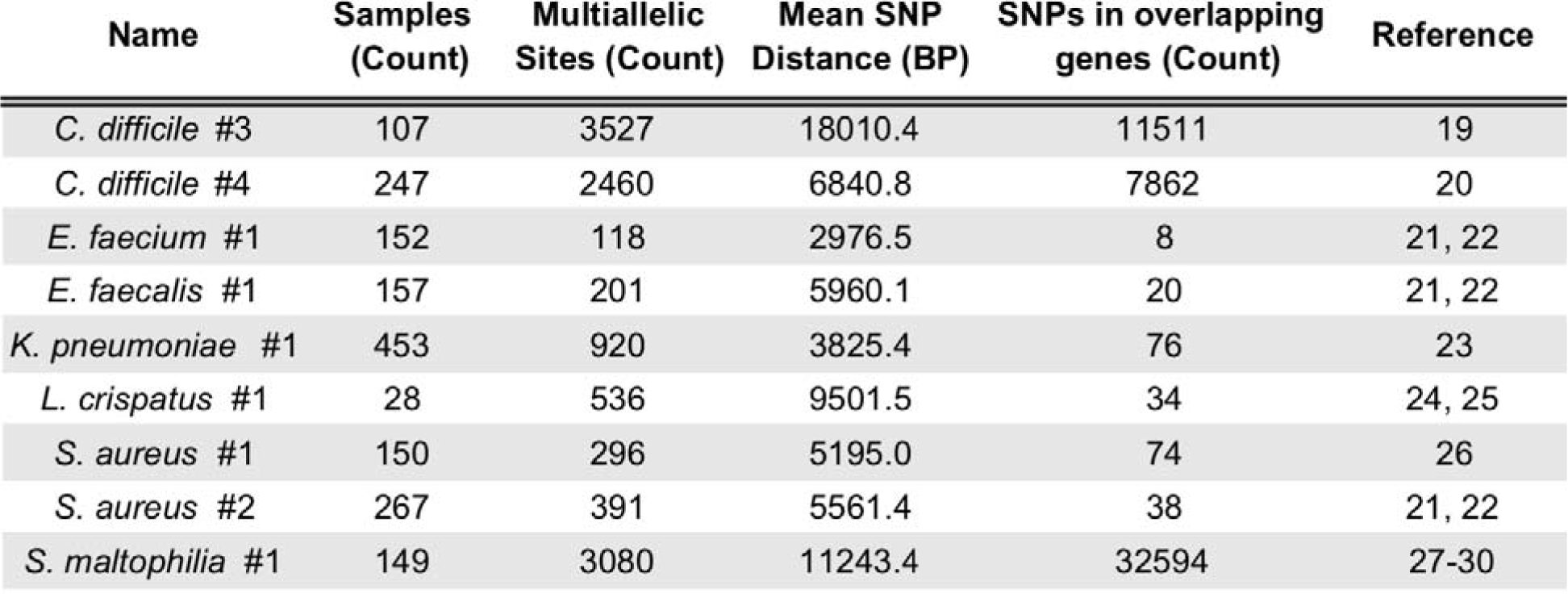
Bacterial datasets.

### Handling multiallelic sites

A multiallelic locus is a site in the genome with more than two alleles present and encompases both triallelic and quadallelic sites. bGWAS typically requires a binary input for each genotype (e.g. 3,4), and multiallelic sites are, by definition, not binary. Thus, special considerations must be taken to use multiallelic sites in bGWAS (see *Multi-line representation for multiallelic sites*). We assessed the potential relevance of multiallelic SNPs to bGWAS on the basis of 1) frequency, 2) differences in functional impact of alternative alleles at a single site, and 3) convergence of multiallelic sites on phylogenetic tree.

#### Multiallelic site frequency

We expected that as the sample size increases the number of multiallelic sites would also increase, as seen across human datasets of different sizes (45); however, this was not the case when looking across different bacterial datasets (Figure S2A). We hypothesized that the lack of correlation between the prevalence of multiallelic sites and dataset size was due to differences in genetic diversity among the datasets (Table 1). Indeed, when we subsample from any single dataset, the fraction of multiallelic sites increases as sample size increases until the diversity of the dataset is exhausted (Figure 2A). Furthermore, datasets with higher sample diversity tend to have a larger fraction of multiallelic sites (Figure 2A, 2B).

**Figure 2.**
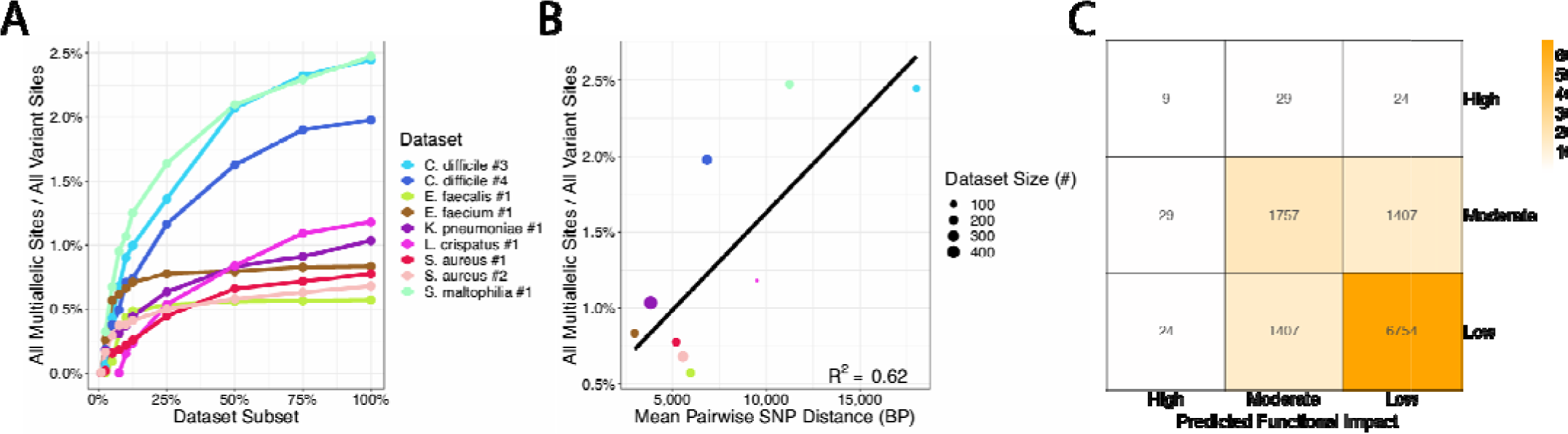
Prevalence and predicted functional impact of multiallelic sites. (A) The number of multiallelic sites increases as sample size increases until the total diversity of the dataset is sampled. (B) More diverse samples have relatively more multiallelic sites. (C) Counts of predicted functional impact (mis)matches for pairs of alleles at triallelic sites (aggregated across all datasets). Alternative alleles often differ in impact.

#### Differences in functional impact

For multiallelic sites, considering each alternative allele at a single site allows for analyses to be performed on alleles based on their predicted functional impact on the encoded protein. Alternative alleles at a single site often have different predicted functional impacts (range across datasets 0-18%, Figure 2C,S1C), and multiallelic sites include alleles with predicted high impact mutations (Figure S2B). In light of these predicted allele-based functional differences, a bGWAS user may want to only run bGWAS on alleles at multiallelic loci that are predicted to have a high impact on the encoded protein.

#### Convergence on phylogenetic tree

For convergence-based bGWAS methods, a significant association between an allele and a phenotype requires that the allele converges on the phylogenetic tree (4,8). If alleles at multiallelic sites are convergent on the phylogeny, then they could potentially contribute to genotype-phenotype associations. We found that single alleles from multiallelic sites are convergent on the phylogeny as often as biallelic sites (Figure S1D), indicating that they could potentially associate with phenotypes when using convergence-based bGWAS.

#### Multi-line representation for multiallelic sites

To use multiallelic sites in bGWAS, these sites typically must be represented as a binary input for each genotype (e.g. 3,4). Three ways multiallelic sites can be handled to fit with the binary framework of bGWAS are: 1) remove them from the dataset prior to analysis, 2) group all minor alleles together, or 3) encode each minor allele separately. Excluding multiallelic sites is problematic if any of these sites determine the phenotype; in these cases, excluding multiallelic sites will result in missed bGWAS hits. Furthermore, coding all minor alleles as one could obscure true associations, particularly if the different minor alleles have dissimilar functional impacts. Multi-line formatting of multiallelic SNPs provides more interpretability, more precise allele classification, and less information loss. For these reasons, multi-line representation is increasingly important in certain human genetics analyses [12] and we propose this same representation for bGWAS studies, particularly for large diverse datasets (Figure 1B).

### Choosing a reference allele

Another aspect to consider when pre-processing SNPs for bGWAS is the allele referencing method, which is critical for a uniform interpretation of variation at a gene locus when grouping SNPs into genes. Three possible allele referencing methods are: the reference genome allele from variant calling, the major allele, or the ancestral allele (Figure 1C). The reference genome allele is the allele found in the reference genome when using a reference genome-based variant calling approach. The major allele is the most common allele at a given locus in the dataset. Neither of these methods encode the alleles with a consistent evolutionary direction. The ancestral allele is the allele inferred to have existed at the most recent common ancestor of the dataset. Given confident ancestral reconstruction, using the ancestral allele as the reference allele allows for a uniform evolutionary interpretation of variants: there is a consistent direction of evolution in that all mutations have arisen over time. We found that the three different methods for identifying the reference allele frequently identify different alleles (range across datasets 0-58%; Figure 3A). Thus, using the reference genome allele or the major allele as the reference allele will not always maintain a consistent direction of evolution for each allele in a gene, obscuring interpretation when grouping variants into genes.

**Figure 3.**
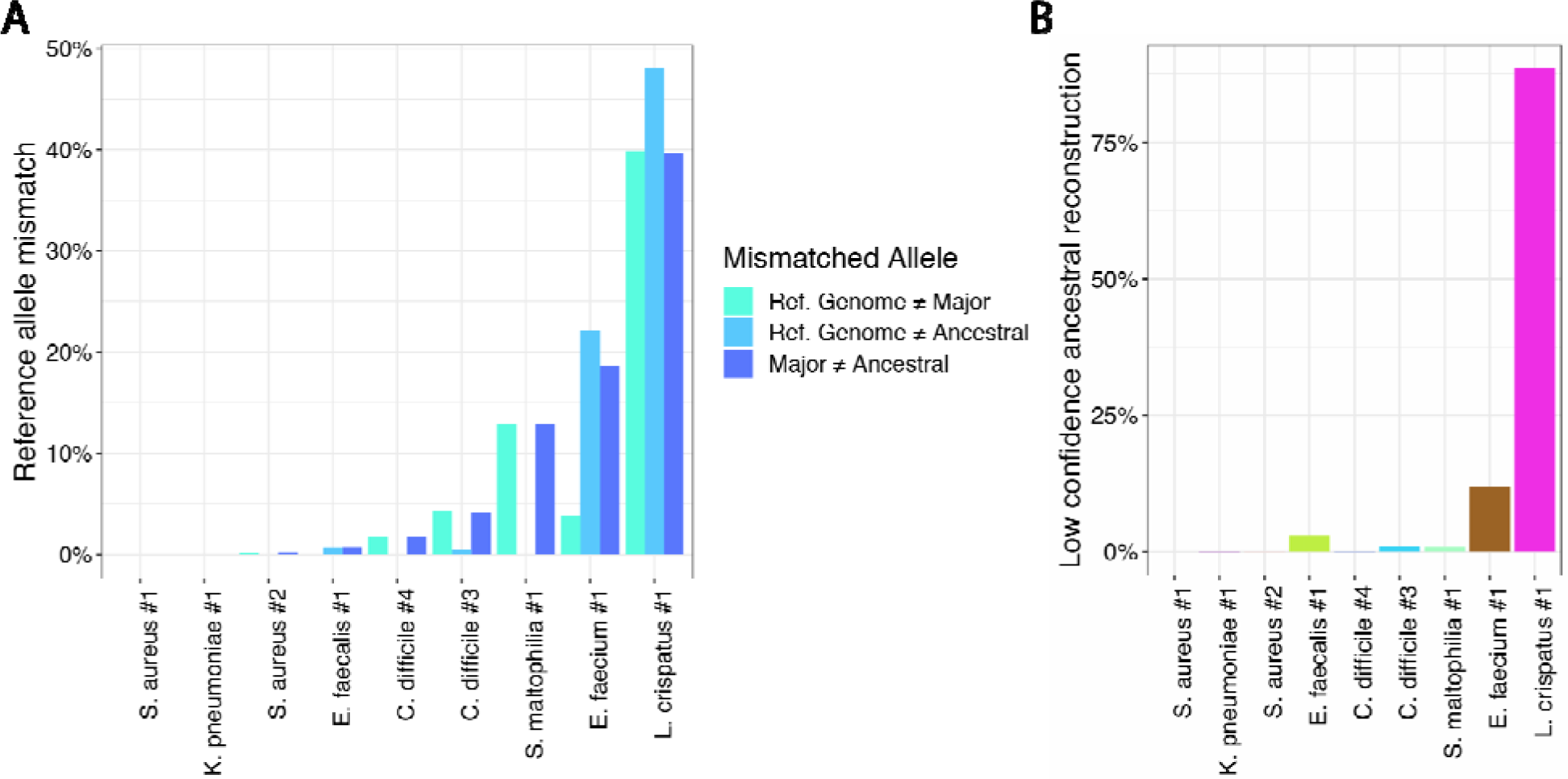
Methods to determine the reference allele identify different alleles. (A) The fraction of variant positions where the identified reference allele varies between two methods. Only high confidence ancestral reconstruction sites (>=87.5% confidence in the ancestral root allele by maximum likelihood) are included. (B) Fraction of low confidence ancestral reconstruction sites for each dataset (<87.5% confidence in the ancestral root allele by maximum likelihood).

Although ancestral reconstruction is the most interpretable option for reference allele choice, this method is not feasible for some datasets. For example, sometimes we cannot confidently predict the most likely ancestral root allele for many loci, as in the *Lactobacillus crispatus* dataset (Figure 3B); in this case, it is not a reliable method to use to define the reference allele. Other limitations of using the ancestral allele as the reference allele are that ancestral reconstruction requires an accurate phylogenetic tree and may be computationally intensive for large datasets. An alternative approach is to use the major allele as the reference allele as this method does not require a tree and thus avoids ancestral reconstruction. When the ancestral allele is not feasible, using the major allele is better than using the reference genome allele when grouping variants into genes because using the major allele leads to less masking of variation at the gene level (Figure S3).

### Grouping variants into genes

Grouping variants into genes prior to performing bGWAS has two advantages for users: 1) improved power to detect genotype-phenotype relationships due to reduced multiple testing burden, and 2) enhanced interpretability as gene function may be clearer than the function of a SNP. Grouping variants into genes may be a particularly helpful approach to bGWAS for datasets with low penetrance of single variants but with convergence at the gene level (Figure 1D). To perform analysis of genomic variants grouped into genes, it is important to consider the choice of reference allele (addressed above), assignment of variants in overlapping genes, and functional impact of the variants.

It is important to ensure that variants in overlapping genes are assigned to each gene that the variant is in to prevent information loss and because the functional impact of a SNP in one gene may be different than its impact on the other gene(s). There are many overlapping genes that share SNPs in each genome (Figure S4A,S4B). Furthermore, there are many sites where the SNP has a different functional impact in the two overlapping genes (cumulative range across datasets 50-70%; Figure 4). The functional impact of variants can be used to select what variants to include in a gene-based analysis. For instance, researchers could subset to only those SNPs most likely to affect gene function (e.g. start loss and stop gain mutations).

**Figure 4:**
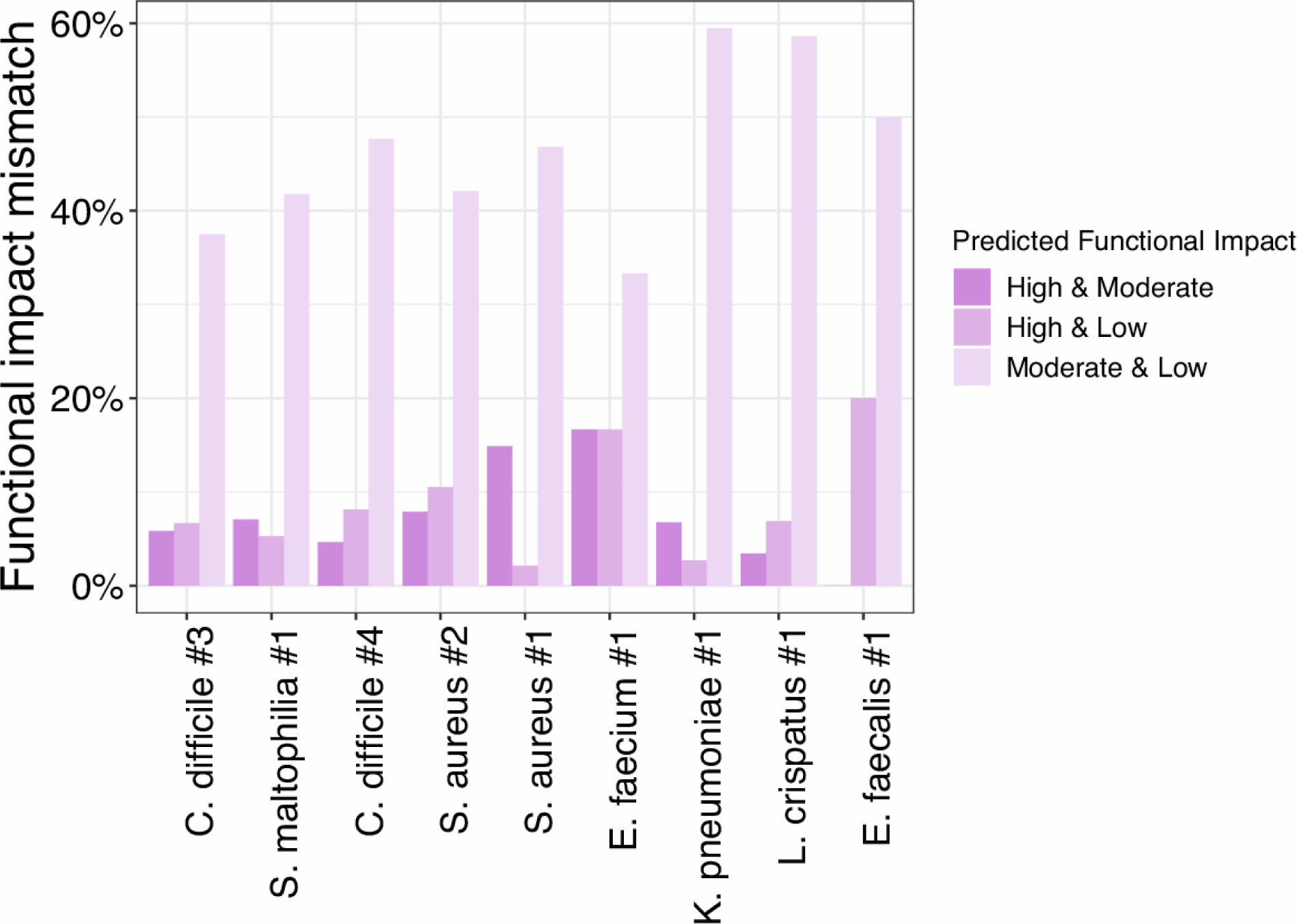
SNPs in overlapping sites can have distinct functional impacts in each gene of the gene pair. The fraction of overlapping variant positions where the SNP has a different predicted functional impact in each of the two overlapping genes.

## PACKAGE DESCRIPTION

We developed prewas to standardize the inclusion and representation of multiallelic sites, choice of reference allele, and SNPs in overlapping genes (Figure 1A) for downstream use in bGWAS analyses. Installation may be performed from GitHub (https://github.com/Snitkin-Lab-Umich/prewas). This R package is an easy-to-use tool with a function that minimally takes a multiVCF input file. The multiVCF encodes the variant nucleotide alleles for all samples. The outputs of the prewas function are matrices of variant presence and absence with multi-line representation of multiallelic sites. Multiple optional files may be used as additional inputs to the prewas function: a phylogenetic tree, an outgroup, and a GFF file. The phylogenetic tree may be added when the user wants to identify ancestral alleles for the allele referencing step. The GFF file contains information on gene location in the reference genome used to call variants and is necessary to generate a binary matrix of presence and absence of variants in each gene. Variants in overlapping genes are assigned to both genes. The matrix outputs from prewas can be directly input into bGWAS tools such as treeWAS (4).

### Generating a binary variant matrix including multiallelic sites (Figure 1B)

The multiVCF file is read into prewas and converted into an allele matrix with single-line representation of each genomic position. Next, a reference allele is chosen for each variant position (see section below). Then, the reference alleles are used to convert the allele matrix into a binary matrix with multi-line representation of each multiallelic site. For each line in the matrix, a 1 represents a single alternate allele, and a 0 represents either the reference allele or any other alternate alleles if the position is a multiallelic site. This binary matrix is output by prewas.

### Identifying reference alleles (Figure 1C)

We have implemented two methods to identify appropriate reference alleles (see Results & Discussion for more details).

#### Ancestral allele approach

The reference allele may be defined as the ancestral allele at each genomic position. In this approach, we identify the most likely allele of the most recent common ancestor of all samples in the dataset by performing ancestral reconstruction. This allele is then always set to 0 in the binary variant matrix. Here, any 1 in the binary variant matrix represents a mutation that has arisen over time, assuming confident ancestral reconstruction results.

#### Major allele approach

The reference allele may also be defined as the major allele at each genomic position. In this case, the most common allele in the dataset is the reference allele. This choice improves the performance speed of prewas as compared to using the ancestral allele at the cost of evolutionary interpretability.

### Grouping variants by gene (Figure 1D)

If a GFF file is provided as input to prewas, variants will be grouped by gene. First, variants found in overlapping genes will be split into multiple lines where each line corresponds to one of the overlapping genes. This ensures that the variant is assigned to each of the genes in which it occurs. Next, variants are collapsed into genes such that the output is a binary matrix with each line corresponding to a single gene and each entry within the matrix is the presence or absence of any variant within that gene.

### Future directions

In a future version of prewas, we plan to implement an option to allow users to select which SNPs they want to include in the binary output matrices based on SnpEff functional impact (e.g. only output predicted high functional impact mutations). When considering the predicted functional impact of each SNP, it is important to use multi-line representation of multiallelic sites even when grouping SNPs by genes because sometimes different alleles at the same site have different predicted functional impacts. Furthermore, prewas could also be extended to process other genomic variants such as indels and structural variants.

## CONCLUSION

We have developed prewas, an easy-to-use R package, that handles multiallelic sites and grouping variants into genes. The prewas package provides a binary SNP matrix output that can be used for SNP-based bGWAS and will prevent the masking of minor alleles during bGWAS analysis. The optional binary gene matrix output can be used for gene-based bGWAS which will enable microbial genomics researchers to maximize the power and interpretability of their bGWAS.

## AUTHOR CONTRIBUTIONS

The study was conceptualized by KS, ZL, SNT, and ESS. Software design and implementation, formal analysis, original draft preparation, and visualization were performed by KS, ZL, and SNT. Data was curated by AP, KS, ZL, and SNT. All authors performed editing and review, and ESS supervised the project.

## ACKNOWLEDGEMENTS

We thank Shawn Hawken for coining the name prewas.

## Data Bibliography

See Table S1.

## SUPPLEMENT

**Supplementary Figure 1:**
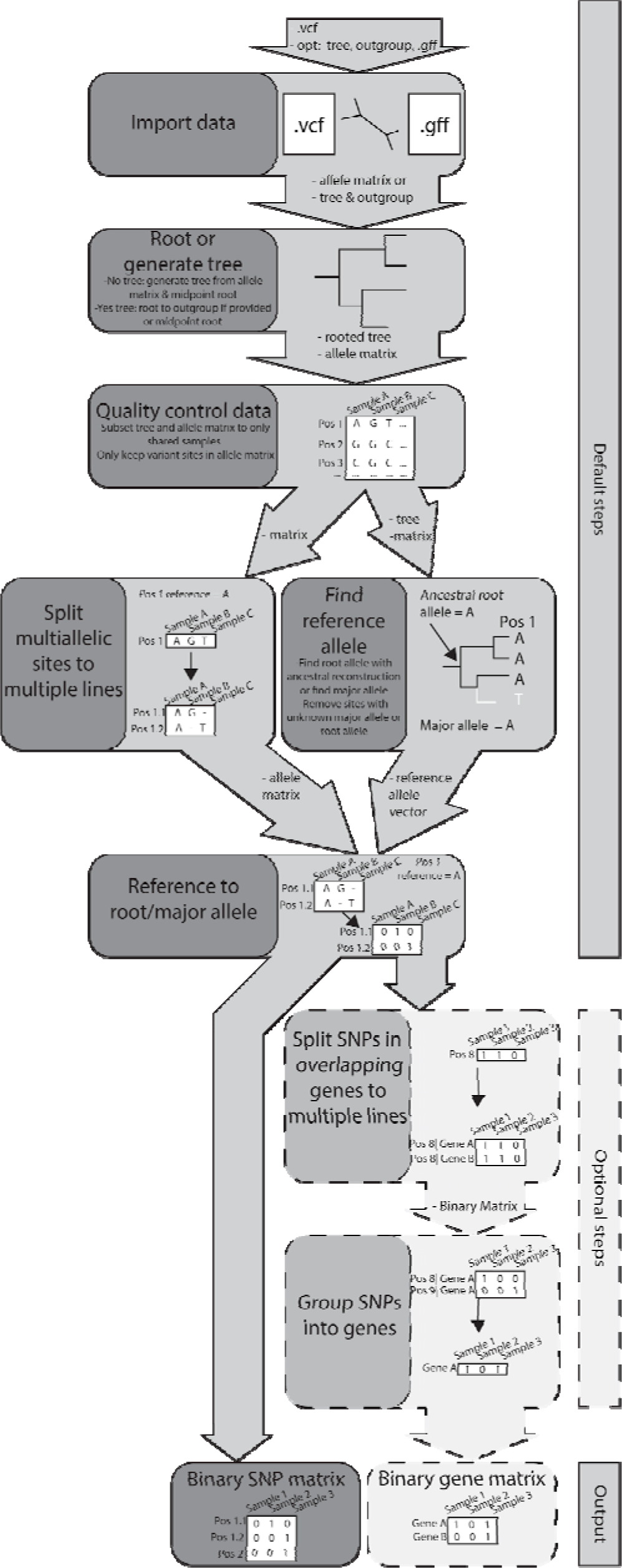
Detailed prewas workflow.

**Supplementary Figure 2:**
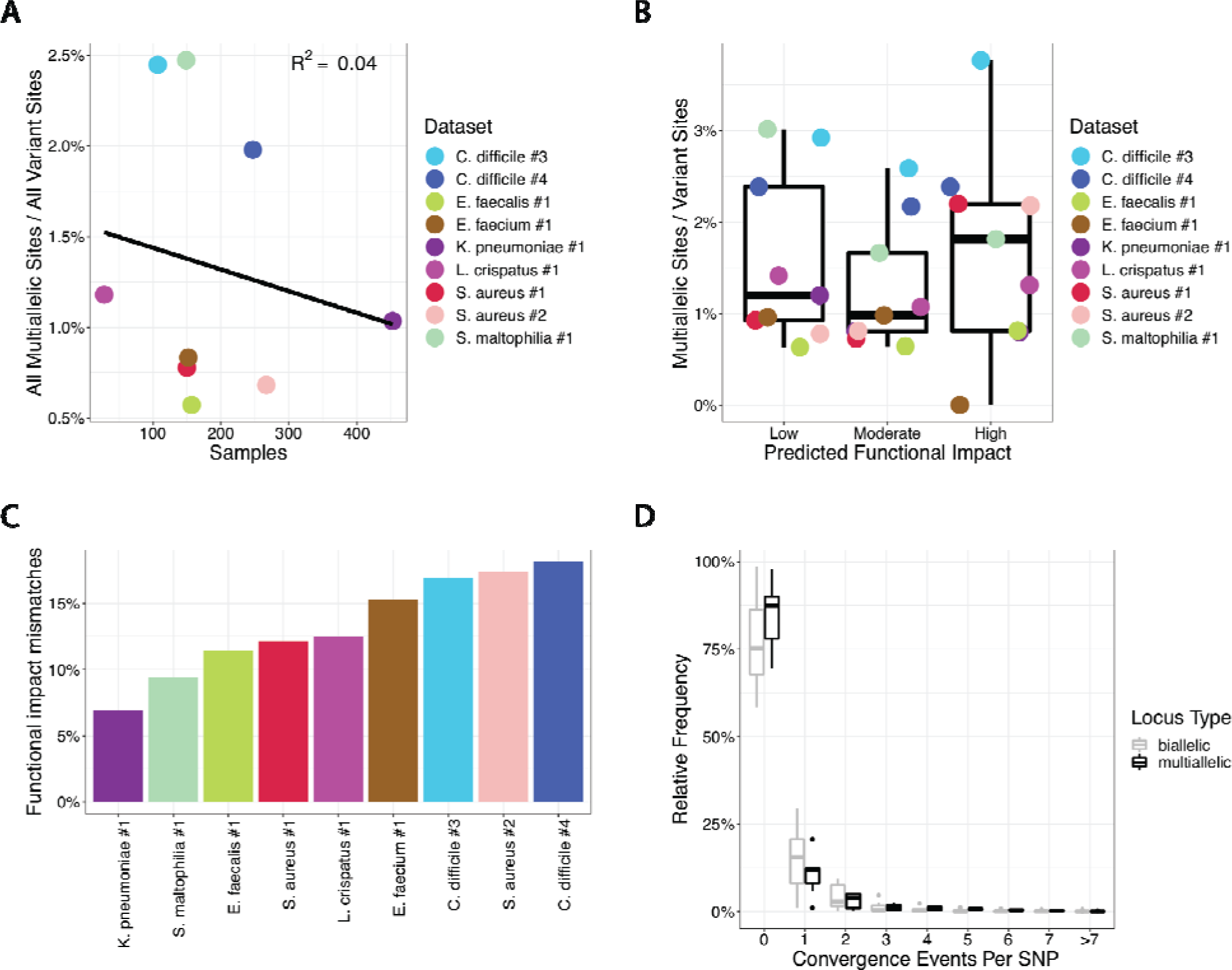
Multiallelic Sites. (A) Independence observed between sample size and prevalence of multiallelic sites. (B) Prevalence of multiallelic sites compared to variant sites with each subset to the various predicted functional impacts. Any multiallelic site with specific impact is compared to any variant site with the same predicted impact. (C) Multiallelic sites with discordant predicted functional impact among alternative alleles. (D) The relative frequency of the number of times an allele arises on the tree. At multiallelic sites, all minor alleles are treated separately.

**Supplementary Figure 3:**
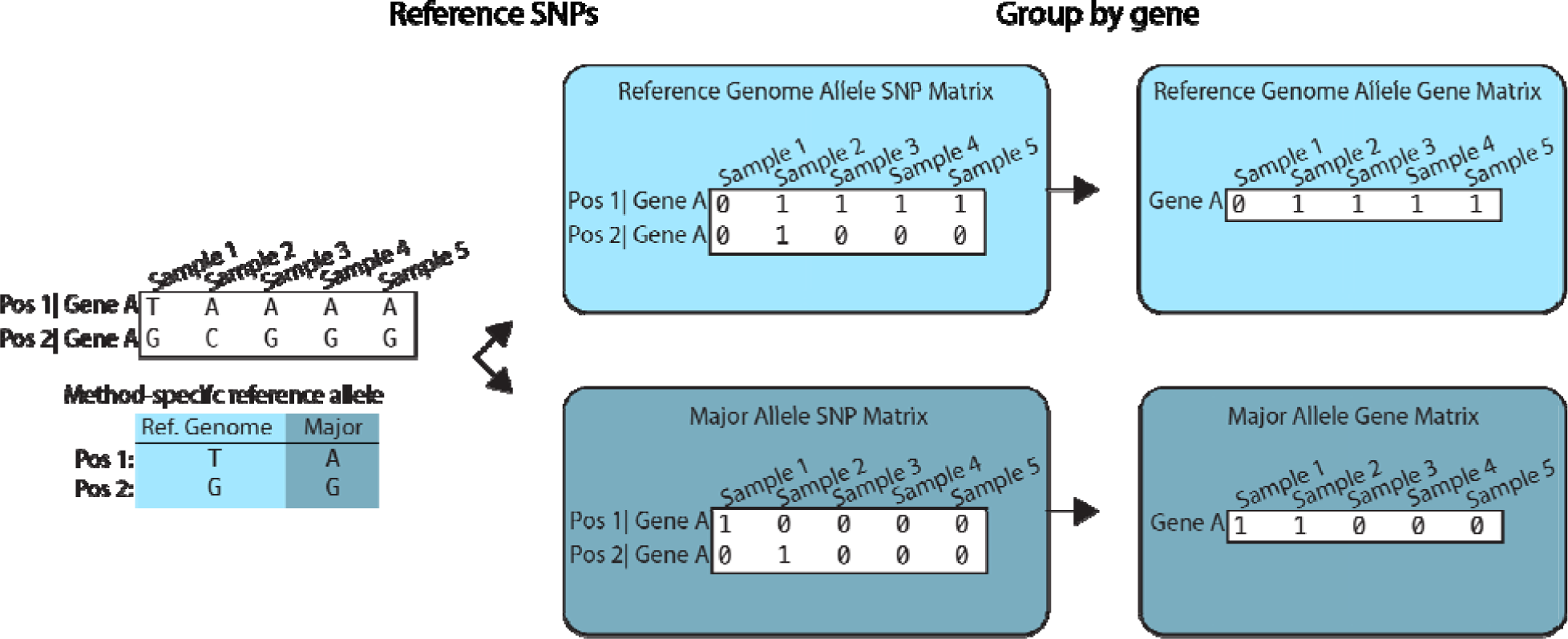
Masking variation at the gene level when grouping into genes. When not confident in the ancestral reconstruction or ancestral reconstruction is not computationally feasible, we suggest referencing to the major allele. In this example, referencing to the reference genome allele masks variation at the gene level. When referencing to the reference genome allele, the variation in Position 2 gets masked by the variation in Position 1 when grouped by gene, leading to a likely lack of association. However, if instead we reference to the major allele, the variation in Gene A is maintained, allowing for potential associations to be detected.

**Supplementary Figure 4.**
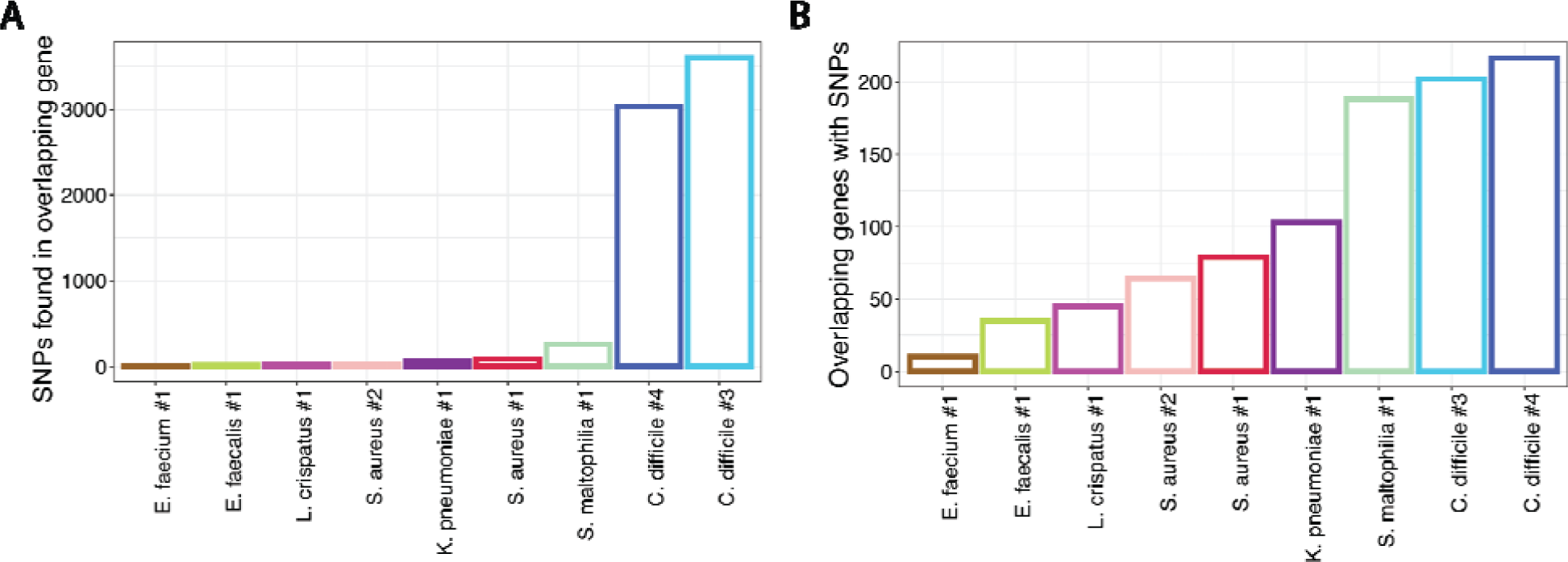
Overlapping genes with SNPs. (A) SNP loci found in positions shared by overlapping genes. (B) Overlapping genes with SNPs found in the overlapping positions.

**Table S1:** Sources for bacterial datasets.

| Name | Dataset Description | Bioproject | Bioproject_link | Reference Genome Biosample | Ref. |
| --- | --- | --- | --- | --- | --- |
| *C. difficile* #3 | Clinical infection isolates | PRJNA594943 | https://www.ncbi.nlm.nih.gov/bioproject/?term=PRJNA594943 | SAMEA1705932 | 19 |
| *C. difficile* #4 | Clinical infection isolates | PRJNA561087 | https://www.ncbi.nlm.nih.gov/bioproject/?term=PRJNA561087 | SAMEA1705932 | 20 |
| *E. faecium* #1 | Healthcare-associated colonization isolates | PRJNA435617 | https://www.ncbi.nlm.nih.gov/bioproject/?term=PRJNA435617 | SAMN10039001 | 21, 22 |
| *E. faecalis* #1 | Healthcare-associated colonization isolates | PRJNA435617 | https://www.ncbi.nlm.nih.gov/bioproject/?term=PRJNA435617 | SAMN10039299 | 21, 22 |
| *K. pneumoniae* #1 | Healthcare-associated clinical isolates | PRJNA415194 | https://www.ncbi.nlm.nih.gov/bioproject/?term=PRJNA415194 | SAMN01057611 | 23 |
| *L. crispatus* #1 | Publicly available genomes | PRJNA547620,<br>PRJNA50051,<br>PRJNA50173,<br>PRJNA50057,<br>PRJNA50067,<br>PRJNA50165,<br>PRJNA50167,<br>PRJNA50053,<br>PRJNA52107,<br>PRJNA52105,<br>PRJNA222257,<br>PRJNA272101,<br>PRJEB8104,<br>PRJNA316969,<br>PRJNA379934,<br>PRJFR22112 | https://www.ncbi.nlm.nih.gov/bioproject/?term=PRJNA547620,<br>https://www.ncbi.nlm.nih.gov/bioproject/?term=PRJNA50051,<br>https://www.ncbi.nlm.nih.gov/bioproject/?term=PRJNA50173,<br>https://www.ncbi.nlm.nih.gov/bioproject/?term=PRJNA50057,<br>https://www.ncbi.nlm.nih.gov/bioproject/?term=PRJNA50067,<br>https://www.ncbi.nlm.nih.gov/bioproject/?term=PRJNA50165,<br>https://www.ncbi.nlm.nih.gov/bioproject/?term=PRJNA50167,<br>https://www.ncbi.nlm.nih.gov/bioproject/?term=PRJNA50053,<br>https://www.ncbi.nlm.nih.gov/bioproject/?term=PRJNA52107,<br>https://www.ncbi.nlm.nih.gov/bioproject/?term=PRJNA52105,<br>https://www.ncbi.nlm.nih.gov/bioproject/?term=PRJNA222257,<br>https://www.ncbi.nlm.nih.gov/bioproject/?term=PRJNA272101,<br>https://www.ncbi.nlm.nih.gov/bioproject/?term=PRJEB8104,<br>https://www.ncbi.nlm.nih.gov/bioproject/?term=PRJNA316969,<br>https://www.ncbi.nlm.nih.gov/bioproject/?term=PRJNA379934,<br>https://www.ncbi.nlm.nih.gov/bioproject/?term=PRJFR22112 | SAMEA2272191 | 24, 25 |
| *S. aureus* #1 | MRSA jail colonization isolates | PRJNA530184 | https://www.ncbi.nlm.nih.gov/bioproject/?term=PRJNA530184 | SAMN00253845 | 26 |
| *S. aureus* #2 | Healthcare-associated colonization isolates | PRJNA435617 | https://www.ncbi.nlm.nih.gov/bioproject/?term=PRJNA435617 | SAMN10038895 | 21, 22 |
| *S. maltophilia* #1 | Publicly available genomes | PRJDB3841,<br>PRJNA267549,<br>PRJNA231221,<br>PRJNA164599,<br>PRJNA380601,<br>PRJNA350620,<br>PRJNA390523,<br>PRJNA483996,<br>PRJNA489399,<br>PRJNA268101,<br>PRJNA344912 | https://www.ncbi.nlm.nih.gov/bioproject/?term=PRJDB3841,<br>https://www.ncbi.nlm.nih.gov/bioproject/?term=PRJNA267549,<br>https://www.ncbi.nlm.nih.gov/bioproject/?term=PRJNA231221,<br>https://www.ncbi.nlm.nih.gov/bioproject/?term=PRJNA164599,<br>https://www.ncbi.nlm.nih.gov/bioproject/?term=PRJNA380601,<br>https://www.ncbi.nlm.nih.gov/bioproject/?term=PRJNA350620,<br>https://www.ncbi.nlm.nih.gov/bioproject/?term=PRJNA390523,<br>https://www.ncbi.nlm.nih.gov/bioproject/?term=PRJNA483996,<br>https://www.ncbi.nlm.nih.gov/bioproject/?term=PRJNA489399,<br>https://www.ncbi.nlm.nih.gov/bioproject/?term=PRJNA268101,<br>https://www.ncbi.nlm.nih.gov/bioproject/?term=PRJNA344912 | SAMEA1705934 | 27-30 |

